# Combining positive and negative regulation for modular and robust biomolecular control architectures

**DOI:** 10.1101/2024.03.22.586143

**Authors:** Kirill Sechkar, Harrsion Steel

## Abstract

Engineered biotechnologies are powered by synthetic gene regulation and control systems, known as genetic circuits, which must be modular and robust to disturbances if they are to perform reliably. An emerging family of regulatory mechanisms is mediated by clustered interspaced palindromic repeats (CRISPR) that can both interfere with (downregulate) or activate (upregulate) a given gene’s expression. However, all CRSIPR regulation relies on a shared resource pool of dCas9 proteins. Hence, a circuit’s components can indirectly affect one another via resource competition – even without any intended interactions between them – which compromises the modularity of synthetic biological designs. Using a resource-aware model of CRISPR regulation, we find that circuit modules which simultaneously subject a gene to CRISPR interference and activation are rendered robust to resource competition crosstalk. Evaluating this architecture’s simulated performance, we identify the scenarios where it can be advantageous over the extant resource competition mitigation strategies. We then consider different feedback architectures to demonstrate that combining opposite regulatory interactions overcomes the trade-off in robustness to perturbations of different nature. The motif of combined positive and negative regulation may therefore give rise to more robust and modular biomolecular controllers, as well as hint at the characteristics of natural systems that possess it.

## 1. Introduction

Synthetic biology aims to engineer cells with desired useful behaviours to tackle industrial, medical and other problems facing humanity. These behaviours involve reacting to different inputs provided to the cell, which it must sense, integrate and act upon in a reliable and predictable manner. To achieve this, cells are conferred with synthetic DNA that encodes genetic circuits, whose genes influence each other’s activity so as to process the information. By combining circuit modules with different functions, highly specific responses to inputs can be achieved [1], [2].

One mechanism that allows genes to affect each other’s expression is mediated by clustered regularly interspaced short palindromic repeats (CRISPR). A single guide RNA (sgRNA) molecule can bind a deactivated Cas9 (dCas9) protein to form a CRISPR complex which recognises and binds with a target DNA sequence within the gene of interest. This can repress the target gene’s activity via CRISPR interference (CRISPRi) [3]. However, recent modifications of the CRISPRi system have allowed to leverage the same mechanism to enable CRISPR activation (CRISPRa) of the target gene instead [1], [4], [5].

Combining sgRNA genes that activate and repress each other or protein-encoding genes (which act as the system’s outputs) can produce a wide range of complex genetic circuits. Since sgRNAs’ targets and effects can be easily altered by changing their sequence, CRISPR circuitry has outstanding flexibility, programmability and multiplexing capacities [4]– [6]. Furthermore, protein production burdens engineered cells, impairing their growth and viability. However, CRISPR circuits only rely on the dCas9 protein, while new functionalities can be added by introducing new sgRNAs, whose synthesis is much less burdensome. CRISPR-based designs can thus be scaled up with little effect on cell growth [6], [7].

However, all CRISPR-mediated interactions depend on the shared pool of free dCas9 proteins. If a given sgRNA’s abundance is increased, it sequesters more of the dCas9 protein, shrinking the overall free resource pool and therefore weakening the regulatory effects of all other sgRNAs in the cell [1], [8]. The possibility of a single circuit element affecting all other CRISPRi/a interactions, even when it is not intended to do so, compromises the modularity of CRISPR circuits [3], [9].

Non-modularity complicates the forecasting of a circuit’s behaviour based on how its components act in isolation. Nonetheless, the performance of CRISPR-based designs can still be predicted and analysed using resource-aware mathematical models [3], [8]. Moreover, by enabling numerical prototyping of genetic circuits, these models allow to develop biomolecular controller architectures aimed at minimising the impact of competition for dCas9 proteins [1], [9].

Expanding the toolkit of such competition-mitigating controllers, in this work we demonstrate that simultaneously subjecting a gene to CRISPR activation and interference renders it robust to fluctuations in resource availability. Using a resource-aware modelling framework, we compare the simulated performances of this and other CRISPR circuit architectures. We then apply the principle of combining CRISPRi and CRISPRa regulation to feedback loop design, showing how it allows to overcome a trade-off in robustness of gene expression to different types of perturbations.

## II. Methods

### A. Mathematical model

We consider the dCas9-AsiA implementation of CRISPRi/a regulation in *E. coli* cells, where a transcription activation factor is directly tethered to the dCas9 protein [4]. Therefore, CRISPRi and CRISPRa complexes are formed via the same mechanism, and the only difference between them is the target DNA sequence’s location in the regulated gene. While CRISPRa complexes land upstream of the gene to let the tethered factor recruit expression machinery at the gene’s promoter, CRISPRi complexes land on or after the promoter to prevent the repressed gene’s DNA sequence from being read [4], [5]. To model this with ordinary differential equations (ODEs), we formulated a framework based on [1] and [9], where sgRNA is assumed to bind and dissociate from dCas9 on a much faster timescale than all other reactions in the cell. The meaning and values of our model’s parameters are given in Table I.

**TABLE I.** PHYSIOLOGICAL MEANING AND VALUES OF MODEL PARAMETERS.

| Parameter | Significance | Value<sup>a</sup> | Units |
| --- | --- | --- | --- |
| $\alpha_{g_i}, \alpha_{g_a}, \alpha_{g_c}$ | Max. sgRNA synthesis rates | $1.5 \cdot 10^4$ | $nM/h$ |
| $\alpha_{g_f}$ | Max. feedback sgRNA synthesis rate | $1.17 \cdot 10^5$ | $nM/h$ |
| $\delta_{g_i}, \delta_{g_a}, \delta_{g_c}, \delta_{g_f}$ | sgRNA degradation rates | 12 | $h^{-1}$ |
| $\lambda$ | Cell growth rate | 0.6 | $h^{-1}$ |
| $K_j$ | sgRNA-dCas9 dissociation constant | 0.01 | $nM$ |
| $Q_j$ | CRISPR complex-target DNA dissociation constant | 0.5 | $nM$ |
| $D_i^T, D_a^T, D_c^T$ | Total target DNA sequence concentration | 200 | $nM$ |
| $D_f^T$ | Total dCas9 gene DNA sequence concentration | 30 | $nM$ |
| $F_a^0$ | Leaky exp. from CRISPR-activatable promoter | 0.05 | None |
| $\alpha_d$ | Max. dCas9 synthesis rate | $1.08 \cdot 10^{3b}$ | $nM/h$ |
| $\alpha_p$ | Max. output protein synthesis rate | $5.04 \cdot 10^3$ | $nM/h$ |

<sup>a</sup>Taken from [4] for leaky expression $F_a^0$ and from [9] for other parameters.
<sup>b</sup> $1.08 \cdot 10^7 nM$ if feedback controller of dCas9 expression is present, since a stronger promoter is used [9].

Let the cell host an output protein gene and some or all of the sgRNA genes from the set *G* = *{i, a, c, f }*. The sgRNAs interfering with and activating the output gene’s expression are labelled *i* and *a* respectively, whereas perturbations to free dCas9 availability are caused by the ‘competitor’ sgRNA *c*. Finally, to benchmark the proposed combined CRISPRi/a architecture against an existing biomolecular feedback controller for mitigating dCas9 competition, we also include a ‘feedback’ sgRNA *f* . The concentration of the unbound sgRNA encoded by a gene *j ∈ G* is governed by (1):

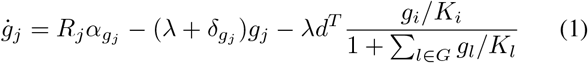

Here, the first term stands for sgRNA synthesis, the second describes its degradation and dilution due to cell division, and the third captures the net removal of sgRNA as it forms CRISPR complexes which then get diluted as the cell dvivides. The variable *d*^*T*^ stands for the total concentration of dCas9 proteins (be they free or bound by sgRNAs), whereas *R*_*j*_ is a circuit-dependent function capturing the regulation of gene *j*’s expression. *R*_*j*_ may thus depend on both CRISPR action and the external input *u*_*j*_, enacted via optogenetic regulation or the addition of chemical inducers to the cell culture medium.

As shown in Fig. 1, the free gene *j* sgRNAs bind dCas9 with an association constant *K*_*j*_ to form CRISPR complexes that have a total concentration of 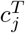 *nM* . Among these, *c*_*j*_ *nM* are free-floating, and the rest are bound to their target DNA sequences with an association constant *Q*_*j*_. This leaves *D*_*j*_ out Of 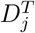 *nM* of target DNA free from CRISPR complexes. All these concentrations can be found according to (2)-(4):

**Fig. 1.**
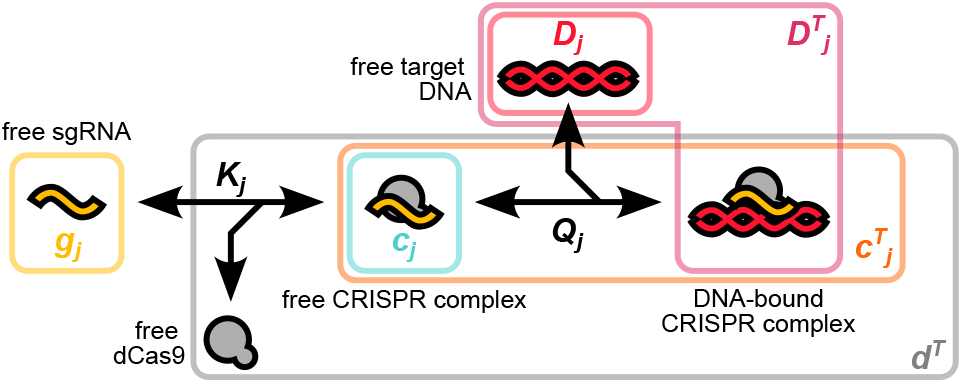
Different states of the sgRNA encoded by a gene *j* and the variables describing their abundances. Note that the total dCas9 level *d*^*T*^ will also include CRISPR complexes with all other sgRNAs if they are present.

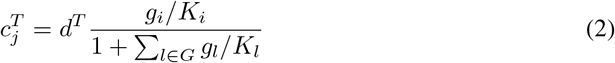

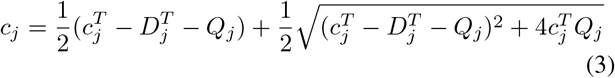

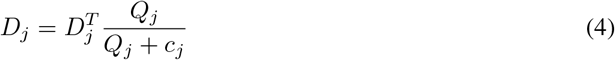

Only the gene copies unhindered by CRISPRi complexes can be expressed. Hence, negative regulation of a target gene by interfering sgRNAs *g*_*i*_ or feedback sgRNAs *g*_*f*_ is captured by the functions *F*_*i*_ and *F*_*f*_, respectively:

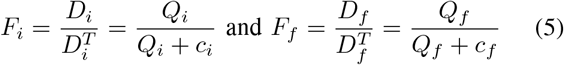

Conversely, while some ‘leaky’ expression of CRISPR-activatable genes without an upstream CRISPRa complex is possible, high expression only occurs when the sgRNA-dCas9 complex is bound to its target. Hence, the effect of CRISPR activation is described by (6).

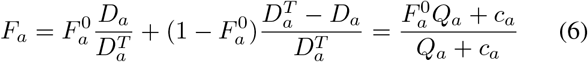

Lastly, since the competing sgRNA is only considered as a disturbance, a definition for *F*_*c*_ is not needed. If a gene is co-regulated by CRISPR interference and activation, the spatially separate locations of target sites allow to assume that CRISPRi and CRISPRa complex binding occurs independently. The overall effect of the two regulatory mechanisms can therefore be found by multiplying *F*_*i*_ by *F*_*a*_ [1].

We also model concentrations of the dCas9 protein (*d*^*T*^) and the circuit’s output protein (*p*), whose dynamics are captured by (7). The positive terms of these ODEs capture protein synthesis and therefore include the circuit-dependent gene regulation functions *R*_*d*_ and *R*_*p*_. The negative terms stand for protein removal by dilution but not active degradation, as the rate of most proteins’ degradation is negligible compared to that of cell growth [10].

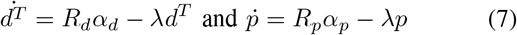

### B. Computational methods

The model, implemented in Python 3.9, was simulated using the diffrax 0.4.0 package’s `Kvaerno3` integrator and the JAX 0.4.12 package. Circuit steady states were retrieved by integrating ODEs over 24 *h* of simulated time. All code used in the study is available at https://github.com/KSechkar/posnegreg.

## III. Results And Discussion

### A. Supplementary repression reinforces positive regulation

Consider a genetic circuit module that consists of a single output protein *p* and an sgRNA *g*_*a*_ that activates its expression (Fig. 2a). Varying the extent of the activating sgRNA gene’s induction *u*_*a*_ (e.g. by adding different chemical inducer concentrations to the cell growth medium) allows to control the output protein’s steady-state level according to the doseresponse curve in Fig. 2c. However, if another sgRNA *g*_*c*_, part of a different circuit module, appears in the same cell, it can sequester dCas9 proteins away from *g*_*a*_, weakening the output gene’s activation. Thus, higher activating sgRNA levels are needed to achieve the same expression of protein, which shifts the dose-response curve. The module of interest’s behaviour is therefore conditional on the rest of the system, which renders the circuit non-modular.

**Fig. 2.**
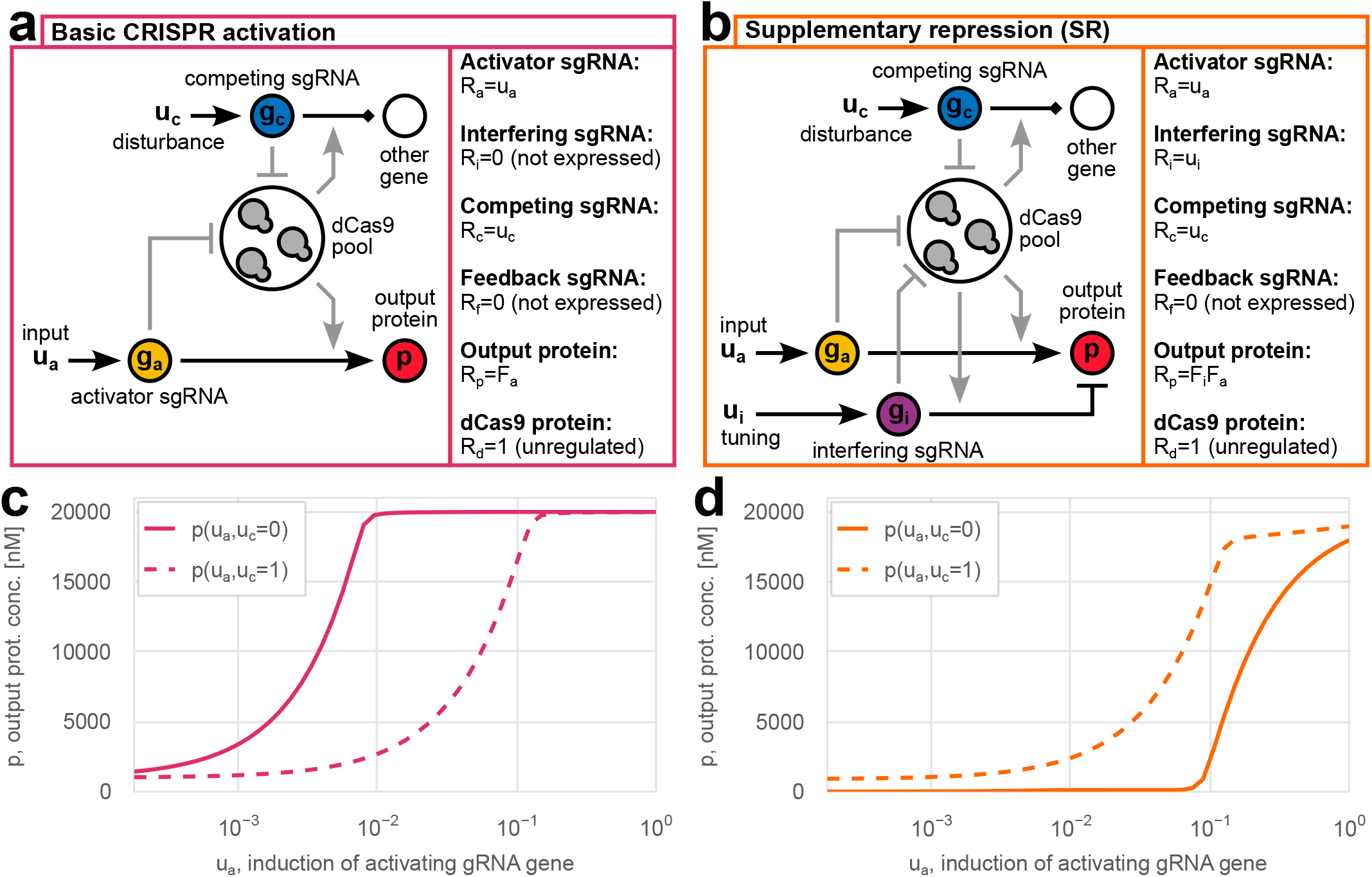
Comparison of basic CRISPR activation and CRISPRa with supplementary repression. (a–b) Schematics and definitions of regulatory functions in (1) and (7) for the two architectures. (c–d) Example dose-response curves with and without the competing sgRNA present for basic CRISPRa and SR, respectively. The SR curve is displayed for *u*_*i*_ = 10^*−*2^.

Let us now augment the module of interest with a supplementary repressive (SR) interaction, enabled by an interfering sgRNA *g*_*i*_, which together with *g*_*a*_ regulates the output gene alongside *g*_*a*_ as shown in Fig. 2b. Then, while the competing sgRNA *g*_*c*_ still reduces resource availability, this weakens both the output gene’s activation by *g*_*a*_ and its repression by *g*_*i*_. The opposite and antagonistic effects of disturbance on the two regulatory interactions mitigate each other, making the doseresponse curve for *u*_*a*_ more consistent with its undisturbed shape and restoring modular behaviour (Fig. 2d). For instance, the half-saturation point 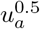 corresponding to *R*_*p*_ = 0.5 changes only *≈* 3-fold with SR instead of *≈* 16-fold in the basic CRISPRa case. Adjusting the interfering sgRNA gene’s induction level *u*_*i*_ allows to tune the strength of supplementary repression to optimise the module’s performance.

Besides the basic CRISPRa and the SR architectures, we also consider the case where *g*_*i*_ is present but acts off-target, regulating not *p* but some other gene (Fig. 3a). This allows to explore the possibility that improved module robustness is caused not by supplementary repression, but rather by a lower baseline resource availability. Indeed, with additional sgRNAs present from the start, the appearance of the same amount of *g*_*c*_ causes a smaller relative change in demand for dCas9. Finally, we benchmark supplementary repression’s ability to mitigate the effects of resource competition against an existing strategy proposed to this end – that is, negative feedback regulation of dCas9 synthesis by a ‘feedback’ sgRNA *g*_*f*_ (Fig. 3b) [9].

**Fig. 3.**
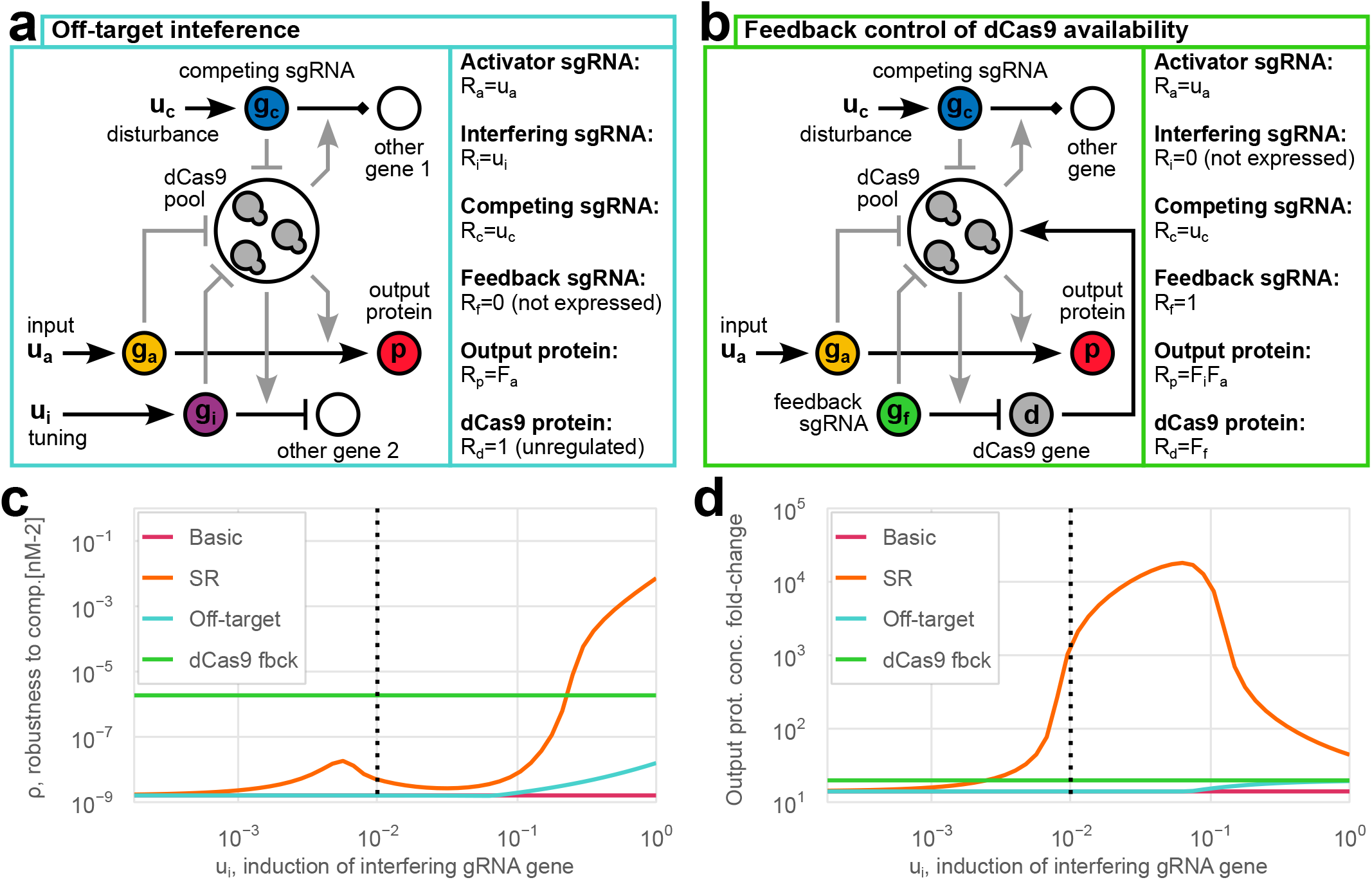
Exploration and benchmarking of the SR architecture performance. (a–b) Schematics and definitions of regulatory functions in (1) and (7) for the off-target interference and feedback control architectures considered in addition to basic CRISPRa and SR from Fig. 2. (c–d) The four considered architectures’ robustness to resource competition *ρ* for different levels of induction of the interfering sgRNA gene *u*_*i*_ between 1.8 *·* 10^*−*4^ (no inducer present) and 1 (full induction). (d) Fold-change in output protein levels between uninduced and fully induced activating sgRNA expression for different *u*_*i*_ values. For the basic CRISPRa and feedback control designs, which do not contain the interfering sgRNA, horizontal lines in (c–d) represent the *u*_*i*_-independent values of the calculated metrics. The dotted vertical line corresponds to the *u*_*i*_ = 10^*−*2^ case displayed in Fig. 2d.

The robustness of all these architectures’ input-output response to competition for dCas9 can be compared using a quantitative metric *ρ* defined in (8). This formula considers the square of the error that the appearance a competing sgRNA (i.e. *u*_*c*_ being set to 1 from 0) introduces to the steady-state output protein level *p* for different extents of the activating sRNA gene’s induction. Integration over the logarithm of *u*_*a*_ mimics serial dilution of a chemical inducer in the culture medium. Meanwhile, the chosen range 1.8 *·* 10^*−*4^ *< u*_*a*_ *<* 1 starts at the extent of leaky expression in absence of any inducer molecules and ends at the full induction of an inducible promoter such as pLux [9].

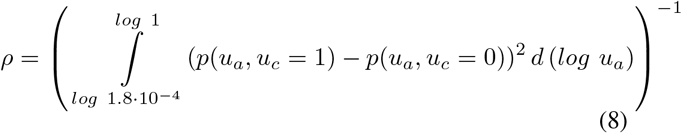

Fig. 3c shows that the SR architecture’s robustness *ρ* exceeds that of both basic CRISPRa regulation and the off-target activation design for all *u*_*i*_ values. Confirming that it mitigates resource competition’s influence on a model, these observations also demonstrate that the performance improvement caused by SR cannot be explained solely by a higher baseline level of competition for CRISPR moieties.

While feedback control of dCas9 expression, conversely, remains more robust to disturbance than the SR design for most *u*_*i*_ values, supplementary repression may be more advantageous for a circuit’s input-responsiveness. Namely, Fig. 3d displays the fold-change in the output protein level as *u*_*a*_ is increased from the minimum of 1.8 *·* 10^*−*4^ to the maximum of 1. When the interfering sgRNA’s expression is sufficiently high (*u*_*i*_ *≥* 2 *·* 10^*−*3^), this fold-change for SR can be several orders of magnitude above that of all other architectures considered, which enables a much clearer contrast between the ‘on’ and ‘off’ states of a module. This improved responsiveness can be explained by the two-fold positive effect of the activating sgRNA on the output. First, there is the direct upregulation of the output gene by CRISPRa complexes. Second, with few activating sgRNAs present, CRISPRi complexes reduce leaky expression by repressing the output gene. When *u*_*a*_ increases, this negative regulation is eliminated as the more abundant activating sgRNAs take dCas9 away from the interfering sgRNA, whose levels remain constant. With the output protein concentration decreased by SR in the ‘off’ state but unaffected in the ‘on’ state, the overall fold-change in the output rises.

### B. Combining negative and positive regulation to overcome the robustness trade-off in feedback control

Besides reinforcing the modularity of open-loop circuit architectures such as our SR design, appreciation of the robustness-improving effect of combined positive and negative regulation can also highlight and overcome the trade-offs in biomolecular feedback controller design.

Negative feedback loops are aimed at achieving reliable and consistent input-output behaviours of a system in the face of external disturbances. Compared to open-loop architectures, systems with feedback control show superior efficiency and robustness to perturbations [11], [12]. On the other hand, feedback loops create a new source of disturbances and variations originating from the circuitry that enables control action [13]. To this kind of disturbances, conversely, open-loop architectures are perfectly robust since they simply lack any of the susceptible elements. The design of reliable and efficient controllers therefore requires achieving a balance between robustness to external and controller-associated perturbations.

For biological systems, external disturbances in this case include shifts in gene expression rates, cell growth rate fluctuations or changes in the cell culturing conditions [11], [12]. A major controller-associated disturbance is competition for the cellular resources required to exert control action, such as dCas9 proteins if feedback is implemented using CRISPR-mediated regulation [2], [3], [9].

To illustrate the trade-off between robustness to these two kinds of disturbance, in Fig. 4 we simulate different architectures’ response to an external transcriptional perturbation and a controller-associated perturbation in the form of increased demand for dCas9 proteins, where smaller peak deviation in the output protein concentration and shorter settling time indicate greater robustness to a given kind of disturbance. Open-loop control of protein levels (Fig. 4a) is indifferent to the appearance of a new sgRNA *g*_*c*_ but highly susceptible to transient changes in the output protein gene’s transcription rate (e.g. due to a temporary shift in the chemical inducer’s concentration). Meanwhile, a simple *cis* feedback loop [12], where the controlled gene represses itself due to being co-expressed with an interfering sgRNA (Fig. 4b), can successfully combat transcriptional perturbations but not controllerassociated disturbances, especially for high *g*_*i*_ production rates. As the *α*_*i*_ value (and thus the feedback strength) falls, the controller’s performance approaches that of the open-loop circuit, becoming resilient to resource perturbations at the cost of robustness to transcriptional disturbances.

**Fig. 4.**
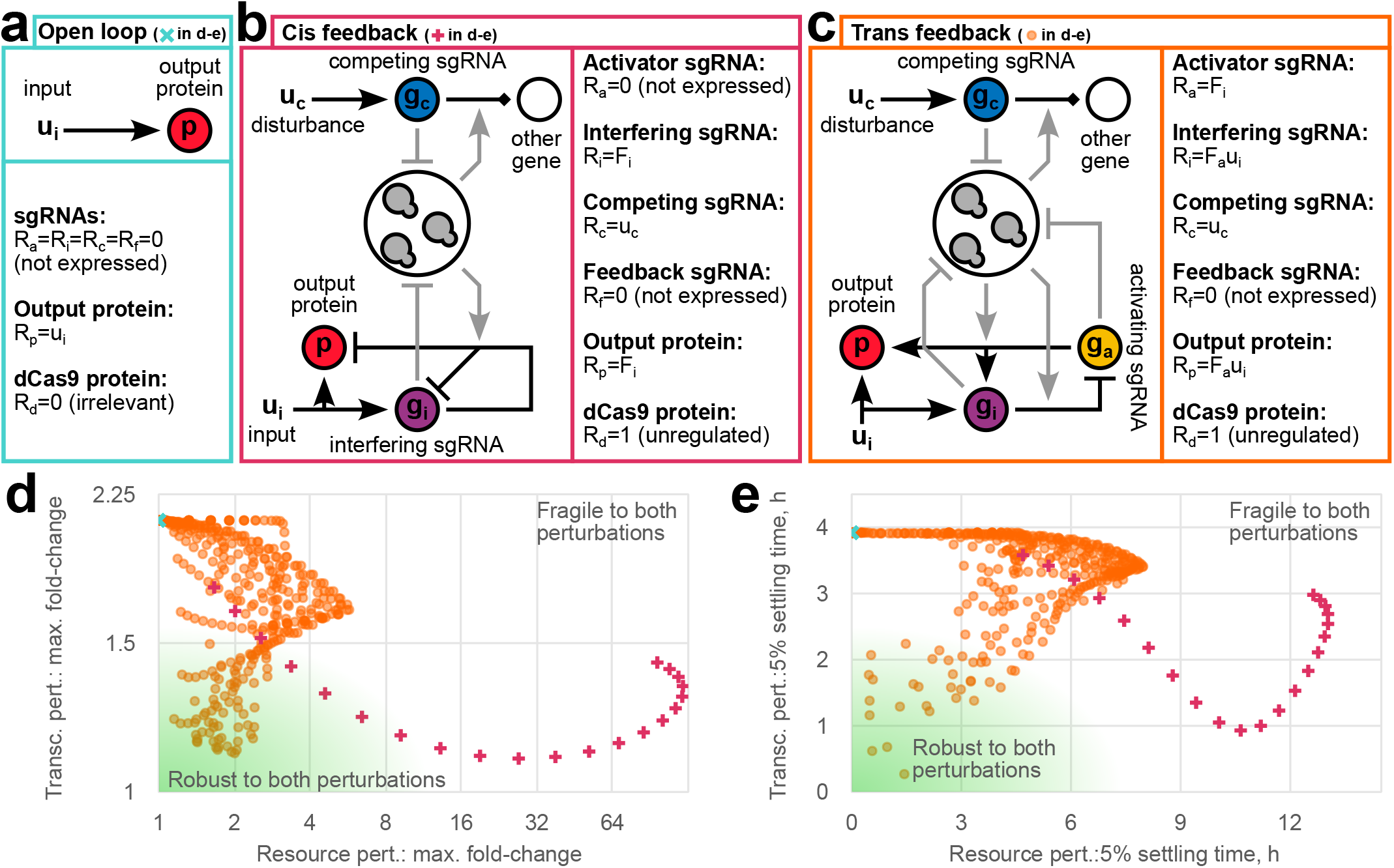
Trade-off between robustness to external and controller-associated perturbations. (a–c) Schematics and definitions of regulatory functions in (1) and (7) for the open-loop, *cis* feedback and *trans* feedback architectures. (d) Maximum fold-changes in the output protein’s concentration in response to external transcriptional perturbation (*u*_*i*_ changed from 1 to 0.25 for 2 hours) and controller-associated resource perturbation (competing sgRNA induced for 2 hours by setting *u*_*c*_ from 0 to 1) for the three architectures with different maximum gRNA synthesis rates. The 21 considered values of *α*_*i*_ were spaced logarithmically between 1.08 *·* 10^2^ *nM/h* and 6 *·* 10^5^ *nM/h*, respectively equivalent to having an uninduced and a fully induced pLux promoter [9]. For *trans* feedback, all combinations of these *α*_*i*_ values with the same 21 possible values of *α*_*a*_ were considered. (e) Time taken by the three architectures considered to return within 5% of the original output protein level after disturbance is removed. The considered perturbations and parameter values are the same as in (d).

However, to reliably maintain the controlled variable at its setpoint regardless of the perturbation, both types of robustness should ideally be maximised. To this end, a controller should be reinforced against the effects of resource competition. Likewise to the previously discussed SR architecture, this can be done by combining CRISPR interference and activation. *Trans* feedback loops include an additional node which is regulated by – and in turn, regulates – the controlled node of interest (Fig. 4c) [12]. Exploring the design space of maximum sgRNA synthesis rates *α*_*i*_ and *α*_*a*_ shows that *trans* feedback occupies the middle between robustness only to external disturbances (strong *cis* feedback) and robustness to just controller-associated perturbations (open-loop system). Moreover, in certain parameter regimes, *trans* feedback can overcome the trade-off between the two kinds of fragility to achieve near-optimal robustness to both disturbances.

### C. Discussion

At its dawn, synthetic biology relied on the assumption of modularity to simplify the process of designing genetic circuits [2], [11]. Since then, crosstalk between different system components via competition for shared resources has been found to be ubiquitous across different biological processes: the expression of all genes in the cell draws from the shared pools of RNA polymerases and ribosomes [14], whereas sgRNAs compete for dCas9 proteins [1], [8], [9]. Meantime, regulation of gene activity by small RNA molecules (sRNAs) can require chaperone proteins from the same pool [15].

At the same time, as more instances of resource competition are discovered, insights from control theory allow to identify common regulatory motifs that help to mitigate the unwanted effects of module crosstalk in different contexts. It likewise helps to determine each of these diverse mechanisms’ strengths and weaknesses, which make it appropriate for particular applications but less suitable for others [2].

A prominent competition-mitigating feature is the combined action of positive and negative regulation that are both affected by changes in the demand for resources. ‘Negative competitive regulation’ allows to mitigate the effects of competition for RNA polymerases and ribosomes enabling gene expression resources by introducing CRISPRi interactions, in which the shared resources, conversely, inhibit expression [16]. Meanwhile, synthetic circuitry can take resources away from the engineered cell’s own genes, slowing down its division, which in turn decreases synthetic protein dilution rate and increases its concentration. To reduce this effect, one can introduce a supplementary repressive element that also experiences up-regulation due to cell growth slowdown, therefore exerting stronger negative regulation on the circuit node of interest [17].

The architectures introduced in this work share the above-mentioned designs’ philosophy of bringing together repression and activation. Moreover, similarly to how it mitigates cell division rate changes, we demonstrate that supplementary repression of a circuit node ensures robustness to changes in the demand for dCas9. On the other hand, combined CRISPRi/a systems possess a property that most previously considered resource competition cases do not [2], [11], [14] – that is, negative and positive regulation being enabled by the same shared resource. This can give rise to unique advantages of designs like the SR architecture discussed in this work, even when alternative resource competition mitigation strategies (like feedback control of dCas9 production) appear more effective. Namely, Fig. 3d demonstrates that the twofold effect of positive regulation in the SR circuit – direct and via the sequestration of resources from repressive complexes – combats leaky expression of the regulated gene, making it more responsive to the user’s input.

These observations’ significance may go beyond CRISPR-mediated circuitry. For instance, regulation by different sRNAs can, too, be both positive and negative while relying on the same shared pool of chaperone proteins [15]. Our findings may thus be applicable to the engineering of genetic circuits relying on these and other shared cellular resources. Furthermore, identifying simultaneous regulation by activation and repression of the same gene in naturally occurring systems may indicate that low leakiness is particularly important for regulatory pathways with such architectures. Meanwhile, the presence of *trans* feedback loops may hint at fluctuations in the availability of resources enabling control action and the resulting need to balance the mitigation of external and controller-associated disturbances.

## IV. Conclusion

In conclusion, this work establishes that the combination of positive and engative regulation promotes modularity and robustness to resource competition in CRISPR-mediated genetic circuits. On one hand, this represents a useful consideration in the design of novel biomolecular controllers, such as the SR circuit proposed here. On the other, it can explain the differences in the behaviour of known feedback architectures like *cis* and *trans* controllers and help to achieve a trade-off between different performance requirements. Given the persistence of resource competition phenomena – and competition mitigation strategies alike – across sundry biological contexts, the insights outlined in this work may be useful in studying and designing other systems with shared resource pools.

## Acknowledgment

K.S. acknowledges support by the Clarendon Fund. H.S. is supported in part by Engineering and Physical Sciences Research Council projects EP/W000326/1 and EP/X017982/1. The authors thank Prof. Y. Schaerli for discussions on the study.

